# *PAPC* couples the Segmentation Clock to somite morphogenesis by regulating N-cadherin dependent adhesion

**DOI:** 10.1101/071084

**Authors:** Jérome Chal, Charlène Guillot, Olivier Pourquié

## Abstract

Vertebrate segmentation is characterized by the periodic formation of epithelial somites from the mesenchymal presomitic mesoderm (PSM). How the rhythmic signaling pulse delivered by the Segmentation Clock is translated into the periodic morphogenesis of somites remains poorly understood. Here, we focused on the role of *Paraxial protocadherin* (*PAPC/Pcdh8*) in this process. We show that in chicken and mouse embryos, PAPC expression is tightly regulated by the Clock and Wavefront system in the posterior PSM. We observed that PAPC exhibits a striking complementary pattern to N-Cadherin (*CDH2*), marking the interface of the future somite boundary in the anterior PSM. Gain and loss of function of *PAPC* in chicken embryos disrupt somite segmentation by altering the CDH2-dependent epithelialization of PSM cells. Our data suggest that clathrin-mediated endocytosis is increased in PAPC expressing cells, subsequently affecting CDH2 internalization in the anterior compartment of the future somite. This in turn generates a differential adhesion interface, allowing formation of the acellular fissure that defines the somite boundary. Thus periodic expression of PAPC downstream of the Segmentation Clock triggers rhythmic endocytosis of CDH2, allowing for segmental de-adhesion and individualization of somites.

## INTRODUCTION

Somitogenesis is an early developmental process whereby pairs of epithelial spheres, called somites, form periodically from the mesenchymal Presomitic Mesoderm (PSM). The periodic arrangement of somites reflects the initial metameric organization of the vertebrate embryo. Somites subsequently differentiate to form the dermis, skeletal muscles and axial skeleton (Chal and Pourquie, 2009). Somitogenesis involves a molecular oscillator, called Segmentation Clock, which drives the periodic expression of cyclic genes and controls coordinated pulses of Notch, FGF and Wnt signaling in the PSM (Hubaud and Pourquie, 2014). These signaling pulses are thought to be translated into the periodic array of somites at a specific level of the PSM called determination front. The determination front is defined as a signaling threshold implemented by posterior gradients of Wnt and FGF (Aulehla et al., 2003; Diez del Corral and Storey, 2004; Dubrulle et al., 2001; Hubaud and Pourquie, 2014; Sawada et al., 2001). Cells of the posterior PSM exhibit mesenchymal characteristics and express Snail-related transcription factors (Dale et al., 2006; Nieto, 2002). In the anterior PSM, cells down-regulate *Snail/Slug* expression and up-regulate epithelialization-promoting factors such as *Paraxis* (Barnes et al., 1997; Sosic et al., 1997). This molecular transition correlates with the anterior PSM cells progressively acquiring epithelial characteristics (Duband et al., 1987; Martins et al., 2009). The first evidence for a segmental pattern is a stripe of expression of the transcription factors of the *Mesoderm posterior 2* (*Mesp2*) family, which are activated at the determination front level downstream of the Clock signal. This stripe defines the position of the future somite boundaries (Morimoto et al., 2005; Oginuma et al., 2008; Saga, 2012). Mesp2 expression becomes subsequently restricted to the rostral compartment of the next somite to form, where its anterior border marks the level of the future somitic boundary (Morimoto et al., 2005).

Somites are generated as a consequence of three key events. The first is the formation of the posterior epithelial wall that bridges the dorsal and ventral epithelial layers of the PSM along the future boundary and allows the formation of the somitic rosette. The second is the formation of an acellular medio-lateral fissure at the level of the future boundary that separates the posterior wall of the forming somite S0 from the anterior PSM (Kulesa and Fraser, 2002; Martins et al., 2009; Watanabe and Takahashi, 2010). The third step consists in the polarization of cells of the somite’s rostral compartment which completes the epithelial rosette formation. Epithelialization of the posterior wall starts before fissure formation at the level of somite S-I (Duband et al., 1987; Pourquie and Tam, 2001; Takahashi et al., 2008). It was shown that *Mesp2* controls the expression of the *EphrinB2* receptor and *EphA4* which promote the epithelialization of the posterior wall of the somite by down-regulating cdc42 activity (Nakajima et al., 2006; Nakaya et al., 2004; Nomura-Kitabayashi et al., 2002; Watanabe et al., 2009). In zebrafish, EphrinB2-EphA4 signaling in the PSM is also implicated in the activation of integrin alpha5 at the forming somite boundary leading to the restriction of fibronectin fibrils assembly in the intersomitic fissure (Koshida et al., 2005). The type I cadherin N-CADHERIN (*CDH2*), which is the major cadherin expressed in the paraxial mesoderm, plays an important role in somite epithelialization and boundary formation (Duband et al., 1987; Horikawa et al., 1999; Linask et al., 1998; Radice et al., 1997). In zebrafish, CDH2 is also required in adjacent PSM cells to maintain integrin alpha5 in an inactive conformation, thus inhibiting the formation of fibronectin fibrils inside the PSM (Julich et al., 2015). This mechanism restricts the production of the fibronectin matrix to the somitic surface and the intersomitic fissure.

*Paraxial protocadherin* (*PAPC/Pcdh8/Arcadlin*) is a protocadherin implicated in paraxial mesoderm segmentation (Kim et al., 2000; Kim et al., 1998; Rhee et al., 2003; Yamamoto et al., 1998), in convergence-extension movements and tissue separation during gastrulation (Hukriede et al., 2003; Kraft et al., 2012; Luu et al., 2015; Medina et al., 2004; Unterseher et al., 2004; Yamamoto et al., 2000), and in synapse remodeling (Yamagata et al., 1999; Yasuda et al., 2007). In the anterior PSM, PAPC is expressed in bilateral stripes under the control of the Notch/Mesp2 signaling pathway (Kim et al., 1998; Rhee et al., 2003). Interfering with PAPC function in the paraxial mesoderm in frog or mouse leads to defects in boundary formation and somite epithelialization (Kim et al., 2000; Rhee et al., 2003; Yamamoto et al., 1998). How PAPC controls somite formation is however not understood.

Here, we performed a molecular analysis of *PAPC* function during somitogenesis in chicken and mouse embryos. We show that segmental expression of PAPC downstream of the Segmentation Clock enhances clathrin-mediated endocytosis dynamics of CDH2, leading to somitic fissure formation through local cell de-adhesion. Thus, PAPC expression stripes in the anterior PSM establish a differential adhesion interface localized at the anterior edge of the PAPC expression domain that delimits the somite boundary.

## MATERIALS AND METHODS

### Embryos

Chicken embryos were staged by days of incubation (e.g., E1, E2), by counting somite pairs and according to Hamburger and Hamilton (HH) (Hamburger, 1992). Wild-type mouse embryos were harvested from timed-mated CD1 mice. *RBPjK-/-*, *Raldh2-/-* and *Wnt3a* hypomorph *Vt/Vt* mutant mouse embryos were obtained by conventional breeding of each line. Embryos were genotyped and phenotyped as described (Greco et al., 1996; Niederreither et al., 1999; Oka et al., 1995). Chicken embryo explants were cultured as described (Delfini et al., 2005) in presence of DAPT [10 μM, Calbiochem](Dale et al., 2003), or SU5402 [80-100 μM, Pfizer, Inc.](Delfini et al., 2005), dissolved in DMSO.

### PAPC isoforms isolation and *In situ* hybridization (ISH)

*PAPC* full length sequence was amplified by PCR from cDNA of E2 chicken embryos. A short (*PAPC-S*; Genbank JN252709) and a long (*PAPC-L*, Genbank EF175382) isoforms were identified. Sequence alignments were done using Vector NTI (Informax). Whole-mount ISH was performed as described (Henrique et al., 1995). Chicken *PAPC* probe was synthesized from ChEST435l18. The chicken *LFNG* (Dale et al., 2003) and the mouse *PAPC* (Rhee et al., 2003) probes have been described. Chicken *N-CADHERIN (CDH2)* probe was amplified by PCR from cDNA using the chicken coding sequence (NM001001615).

### PAPC antibodies generation and Western blot analyses

cDNA coding for fragments of the extracellular domain of chicken PAPC and mouse PAPC were cloned in pET vector expression system (Novagen), expressed in *E.coli*, purified with His-Bind Kit (Novagen) and used to immunize rabbits (Cocalico Biologicals, Inc.). The sera were collected, assayed and validated by Western blot and used as anti-PAPC polyclonal antibodies.

Western blot analysis was done following standard procedures. Protein extracts were obtained by lysis in RIPA buffer of pools of dissected tissue of E2 to E3 chicken embryos after electroporation. The signal was detected with an HRP-conjugated anti-rabbit IgG (1:1,000) and an ECL+ kit (Amersham).

### Plasmids and *in ovo* electroporation

*In ovo* electroporations were performed as described (Delfini et al., 2005). Full-length coding sequences of chicken *MESO2* (Buchberger et al., 2002) and *MESPO* (Buchberger et al., 2000) were cloned in pCIG (Megason and McMahon, 2002). The coding sequence of *PAPC-S* was subcloned in pCImG (pCAAGS-IRES-membrane GFP). PAPC RNAi targeting sequences were designed using Genscript and cloned into the RNA interference vector pRFP-RNAi containing a RFP reporter (Das et al., 2006). pBIC (pBI-Cherry) was generated from the Tet-inducible bi-directional promoter pBI (Clontech) by subcloning mCherry in pBI. pBIC-*CDH2* (pBICN) was then generated by subcloning the coding sequence of chicken *CDH2* into pBIC. pBIC-*CDH2* was co-electroporated with pCAGGS-rtTA, and Doxycycline (1μg/mL) was added *in ovo* after overnight incubation and embryos were reincubated further 7-10 hours before fixation. Control embryos were electroporated with matched empty vectors, namely pCIG, pCImG, pRFP or pBIC. After electroporation, embryos were reincubated for 25-30 hours. Embryos were then fixed and fluorescent reporter expression was analyzed before ISH and immuno-histochemistry processing.

### Immunohistochemistry and electron microscopy

Whole-mount immunohistochemistry was performed essentially as described (Bessho et al., 2003). Embryos were incubated with anti-chicken PAPC (1:8,000) anti-mouse PAPC (1:8,000) and anti-CDH2 (Sigma; 1:1,000) at 4°C for two days. Secondary antibodies conjugated either with HRP or AlexaFluor (Molecular probes) were used at 1:1,000. For cryosections (12μm) immunolabeling, working dilution of the anti-chicken PAPC was 1:2,000, anti-CDH2 was 1:300, anti-Clathrin (Cell signaling) was 1:300, anti-ZO-1 (Zymed) was 1:50 and anti-GFP (Abcam) was 1:1,000. Antibodies were incubated at 4°C overnight. F-Actin was detected with fluorescent Phalloidin (ThermoFisher Scientific) used at 1:300 and incubated at 4°C overnight. Samples were then analyzed by confocal microscopy with a Zeiss LSM5 Pascal, LSM780 or Leica SP2. For electron microscopy analysis, explants composed of the PSM, last formed somite and the associated neural tube were dissected and fixed for 1 hour at room temperature in 4% paraformaldehyde (PFA) with 2.5% glutaraldehyde in 0.1 M PBS. Following fixation, tissue was prepared for ultrathin (60 nm) frontal sections and stained for EM analysis. For immunolabelling, PSM was fixed for 1 hour at room temperature in 2% paraformaldehyde with 0.01% glutaraldehyde in 0.1 M PBS, 80 nm sections were incubated with anti-PAPC (1:200) and secondary antibody conjugated to gold beads (10 nm) at room temperature for 1 hour (Amersham, Piscataway, NJ). Sections were post-stained in uranyl acetate. Analysis was performed on a FEI microscope at 80 kV.

### PAPC-CDH2 Colocalization measurement

Signal intensity and distribution for CDH2, PAPC and F-ACTIN (Phalloidin) on immunostained parasagittal chicken embryo sections were analyzed both at the tissue level (PSM areas) and subcellular level (cell-cell junctions).

PSM area level colocalization analysis was performed in Image J using the Coloc2 plugin. Each PSM subdomains was divided in 5 areas of 19μm^2^ and used for subsequent quantification (n=3 embryos, 15 squares per subdomains). For each PSM area, the signal intensity and distribution for CDH2, PAPC and F-Actin stainings were compared 2 by 2 and a Pearson’s coefficient was calculated (ranging from −1 to 1, with 1 corresponding to a total positive correlation). The analysis was done at a 200nm resolution.

Junction level co-localization analysis was performed in Image J. Along each cell-cell junction, the mean junctional signal intensity was collected using Plot profile function (line width 3 pixels) in Image J. Each signal was then cross-correlated 2 by 2 using IGORPro software (macro from (Munjal et al., 2015)) which generate a Pearson’s coefficient for each pixel. A peak at 0 micron means that both signals are co-localized. Statistical significance was assessed using Kolmogorov-Smirnov test (n=20 junctions).

### Phenotype quantifications

The distribution of electroporated cells was quantified on confocal images of parasagittal sections of the last three formed somites. Data were collected for at least six embryos per condition, with two to three sections per embryo analyzed. R-C distribution: distribution of electroporated cells between the rostral versus the caudal halves of newly formed somites. The mesenchymal index was defined as the ratio between the mesenchymal versus epithelial fraction of the cells by direct scoring of cells’ location and morphology. Epithelial cells were defined as cells within the epithelial ring and exhibiting centripetal polarization based on F-Actin, CDH2 and fluorescent reporter expressions. Mesenchymal cells were defined as non polarized cells within the somitocoele, the epithelial ring, and cells outside the ring structure when present. Statistical significance was assessed using Student’s t-test.

For the cell-cell connectivity index, the anterior PSM subdomains S-I and S0 were first identified at low magnification by the overall tissue morphology and the presence of a forming acellular fissure. Then four high magnification 200μm^2^ micrographs for each subdomains were acquired. Individual cell-cell contacts (number and length) were quantified using Image J. The tissue cell-cell connectivity index was defined as the average length of cell-cell contact per cell, data are represented normalized to control S0 caudal domain value, fixed at 100. Electroporated embryos were processed in parallel. Two embryos per condition were analyzed. Statistical significance was assessed using Student’s t-test.

### Endocytosis assays

Chicken embryos were electroporated with pCImG-PAPC-S or pCImG at stage 5HH then cultured on filter paper on agar/albumen plate (Chapman et al., 2001) for 24 hours. After 24 hours, embryos were treated for 20 min with a single 10μL drop of DMSO or of the clathrin- mediated endocytosis inhibitor Pitstop2 at 30 μM (Abcam, ab120687) deposited on the embryo’s ventral side. Next, a sagittal slit was generated within the PSM using a tungsten needle and embryos were incubated with Dextran for 7 min at 37C (2μl of a 1mg/mL Dextran -PBS solution; 10,000MW, conjugated with Alexa Fluor 647; Molecular probes). Next, embryos were washed in cold PBS for 2 min at 4C and fixed in PFA 4% overnight for further immuno-staining processing. Dextran uptake by electroporated cells was measured as the intensity of the retained Dextran fluorescent signal after washes of the treated embryos. Number of embryos analyzed per conditions: pCImG/DMSO n=2, pCImG/Pitsop2 n=4, PAPC-S/DMSO n=7, PAPC-S/Pitstop2 n=6. For each embryos, electroporated PSM was subdivided in ∼5 regions and corresponding Dextran signal intensity in GFP positive versus GFP negative cells was measured using Image J ( >110 cells per conditions). For Chlorpromazine treatment, bisected posterior embryo explants were cultured for 3-5 hours as described (Delfini et al., 2005), left side treated with DMSO (control) and right side with Chlorpromazine at 50 μM (Sigma). Three embryos per conditions were analyzed.

## RESULTS

### *PAPC* (*Pcdh8*) expression domain defines the future somitic boundary

We isolated two distinct, full-length PAPC coding sequences from chicken embryo cDNA (Accession number EF175382 and JN252709), resulting from the differential splicing of the 3’ end of exon1 (Fig. 1A). Both isoforms code for transmembrane proteins composed of an extracellular domain including six Extracellular Cadherin (EC) motifs, a single transmembrane domain and an intracytoplasmic tail (Fig. 1A). The PAPC short isoform (*PAPC-S*) is lacking a 47 amino-acid stretch in its cytoplasmic domain, compared to the long isoform (*PAPC-L*, blue domain) (Fig. 1A). These two isoforms are similar to that described in mouse (Makarenkova et al., 2005). We next generated a polyclonal antibody against the extracellular domain of the chicken PAPC proteins. In PSM protein extracts, PAPC appears as a doublet around 110kD, close to the predicted molecular weight of the isoforms, 103 and 108 kD, respectively, with the long isoform appearing more abundant (Fig. 1B).

**Fig. 1.**
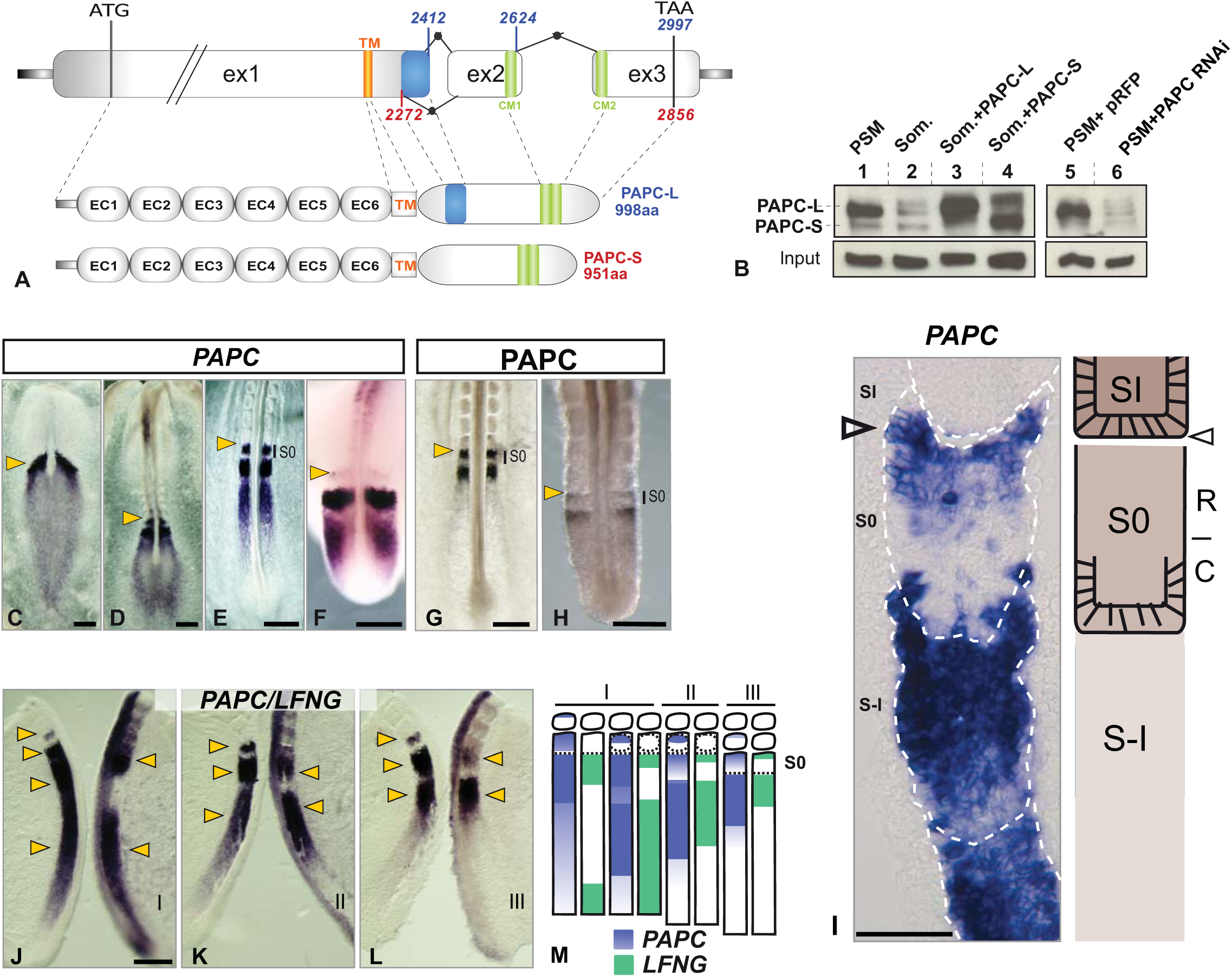
Characterization of chicken *Paraxial protocadherin*. **(A)** The *PAPC* locus is composed of three exons (ex1 to ex3). Sequence features are indicated (in base pairs). The long (*PAPC-L*) and short (*PAPC-S*) isoforms differ by the alternative splicing of 140 base pairs at the 3’ end of exon1 (blue box). EC: Extracellular cadherin motif; TM: transmembrane domain; CM1/2: conserved domains of δ-protocadherins (green boxes). **(B)** Western blot protein analysis using the chicken PAPC antibody in extracts of wild-type PSM (lane1), wild-type Somite (2), somites overexpressing *PAPC-L* (3), *PAPC-S* isoform (4), and PSM expressing *PAPC* RNAi constructs (5,6). **(C-H)** Expression of *PAPC* mRNA in chicken embryo at stage 6HH (C), 6-somite stage (D), E2 (20-somite) embryo (E) E3 embryo (F), and of PAPC protein in E2 (20-somite) embryo (G), and in mouse at E10.5 dpc (H). Whole embryo (C-D) and detail of the posterior region showing the PSM (E-H). S0: forming somite. Arrowheads denote last formed somite boundary. **(I)** (left) Parasagittal section showing chicken *PAPC* mRNA expression in the anterior PSM (blue). Anterior to the top. Somite boundaries are delimited by white dashed lines. (right) Corresponding diagram, S-I/0/I:somite -I/0/I; R:rostral; C:caudal. **(J-M)** Direct comparison of *PAPC* and *LFNG* mRNA dynamic expression on bissected E2 (20-somite) chicken embryos (J-L) and schematic comparison of *PAPC* and *LFNG* expression during the formation of one somite (M). Arrowheads denote expression stripes. Phases of the Segmentation Clock cycle are indicated by roman numerals. (C-H, J-L) Dorsal views, anterior to the top. Scale bar, (C-L) 200μm, (I) 50μm.

During chicken embryo development, *PAPC* mRNA expression is first detected at stage 4HH in the newly ingressed paraxial mesoderm (data not shown). From the onset of somitogenesis (stage 7HH; day 1) to the end (stage 24HH; day 4), *PAPC* expression formed a marked, decreasing rostrocaudal gradient along the PSM, with two or three stripes in the anterior PSM (Fig. 1C-F). *PAPC* mRNA expression is observed by *in situ* hybridization in the posterior PSM, while no protein could be detected (Compare Fig. 1E and G). In the anterior-most PSM and forming somite (S-I to SI), *PAPC* mRNA and protein were detected in a striped pattern (Fig. 1E-H). In the anterior PSM, *PAPC* mRNA expression becomes restricted to the rostral compartment of the forming somite, creating an interface at the level of the forming boundary (Fig.1I).

### The dynamic expression of PAPC is downstream of the Segmentation Clock

We noticed different *PAPC* expression patterns in the PSM of chicken embryos with exactly the same somite number, suggesting that *PAPC* expression is highly dynamic (Fig.1J-L). In some embryos, *PAPC* expression extended along the posterior PSM; whereas in others, it was restricted to the anterior PSM (Fig. 1J-L). Direct comparison of *PAPC* expression with the cyclic gene *LUNATIC FRINGE* (*LFNG*) (McGrew et al., 1998), shows that *PAPC* expression also follows a periodic sequence (Fig.1J-M, n=17). In contrast to *LFNG*, however, *PAPC* is never detected in the most caudal part of the PSM. During phase I of the *LFNG* cycle (Pourquie and Tam, 2001), when *LFNG* is expressed in a broad posterior domain, *PAPC* is also expressed in a broad, gradient-like domain in the posterior PSM (Fig. 1J). As the *LFNG* expression domain moves and narrows anteriorly, the *PAPC* expression domain likewise becomes restricted to the anterior PSM (Fig.1K-M). This dynamic expression in the posterior PSM suggests that *PAPC* defines a new class of cyclic gene regulated by the Segmentation Clock (Fig.1M).

Since the segmentation clock is mainly regulated by FGF, Wnt and Notch signaling we next explored the role of these signaling pathways in regulating *PAPC* dynamic expression. Strikingly, compared to FGF signaling, *PAPC* exhibits a reverse expression gradient (decreasing caudally) in the PSM. We tested whether *PAPC* expression in the posterior PSM is regulated by FGF signaling, using SU5402, a FGF signaling inhibitor (Mohammadi et al., 1997). In treated embryos, *PAPC* expression was strongly up-regulated throughout the PSM compared to control DMSO-treated embryos (Fig. 2A, B; n=7) indicating that FGF signaling represses *PAPC* expression in the posterior PSM. *Mesogenin1*, a transcription factor expressed in the posterior PSM downstream of Wnt signaling, plays a key role in paraxial mesoderm patterning in the mouse embryo (Buchberger et al., 2000; Wittler et al., 2007; Yoon and Wold, 2000). Overexpression by electroporation of the chicken *Mesogenin1* homologue called *MESPO* (Buchberger et al., 2000) in the PSM resulted in strong ectopic expression of *PAPC* throughout the paraxial mesoderm, except in the most caudal region (Fig. 2E, F; n=9). These data suggest that a periodic FGF input inhibits the Wnt/Mesogenin-dependent *PAPC* expression in the posterior PSM, resulting in the cyclic transcription of *PAPC*.

**Fig. 2.**
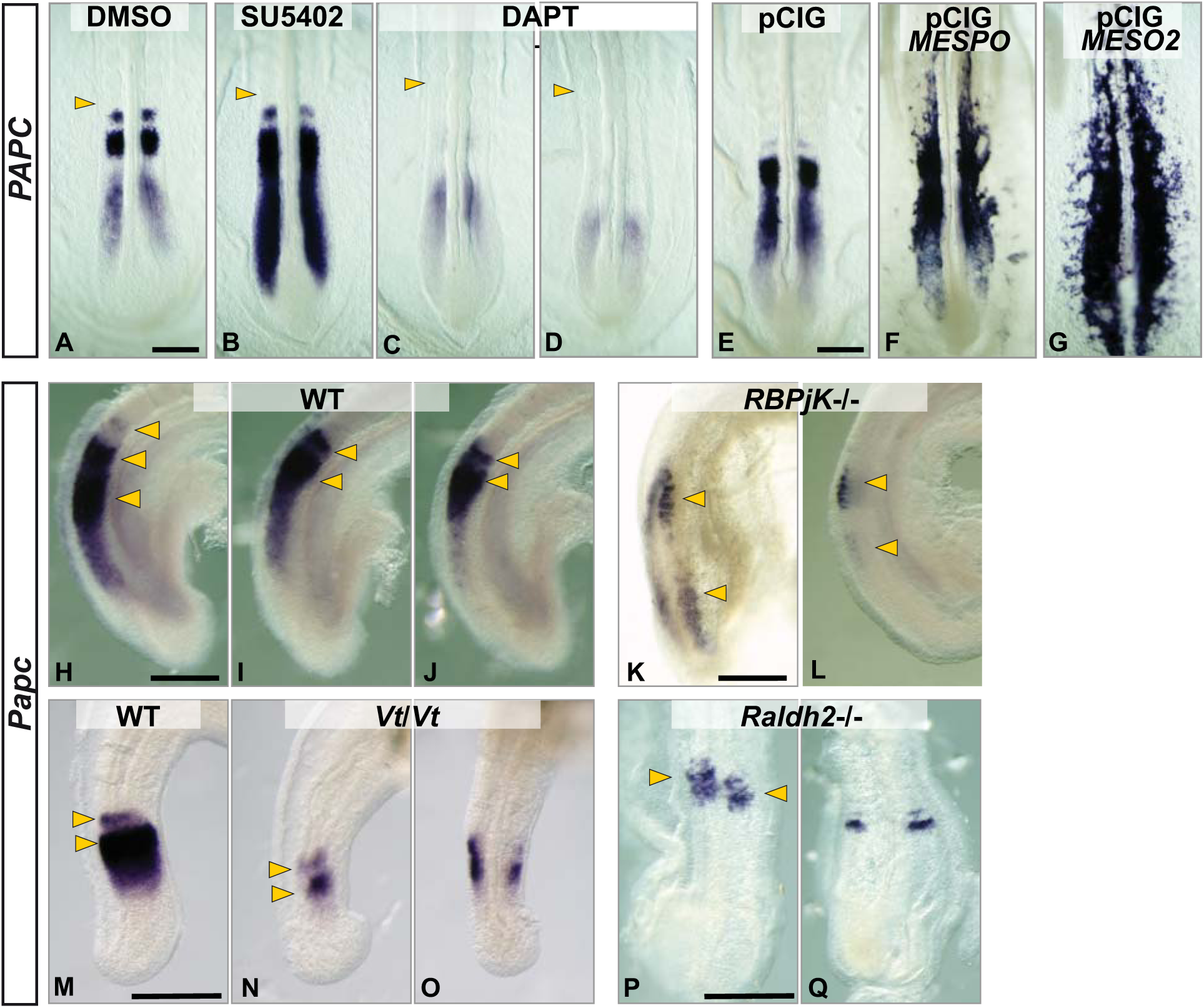
The periodic expression of *PAPC* is controled by FGF/Wnt signaling in the posterior PSM and by Notch/Mesp2 in the anterior PSM. **(A-D)** Whole mount *PAPC* in situ hybridization of E2 chicken embryo posterior explants cultured for 3-4 hours in the presence of DMSO (0.2%) (A), SU5402 (80μM) (B) and DAPT (10μM) (C,D). Arrowheads indicate the last formed somite. **(E-G)** Whole mount *PAPC* in situ hybridization of E2 chicken embryos over-expressing a pCIG control vector (E) a *MESPO* (F) or a *MESO2* (G) expressing vector in the PSM. **(H-J)** Whole mount in situ hybridization showing periodic expression of the mouse *Papc* RNA in the PSM of stage-matched E9.5dpc wild-type (WT) embryos. Arrowheads indicate the anterior boundary of expression stripes. **(K-Q)** Whole mount in situ hybridization showing Papc expression in mice mutant for *Rbp-jK*-/- (E9.0dpc) (K, L); in control (M) and *Vestigial tail* mutants (*Vt/Vt*) (E10.5dpc) (N–O) and in *Raldh2*-/- mutants (E8.5dpc) (P, Q). Arrowheads indicate expression stripes. Anterior to the top, lateral view (H-N), dorsal view (A-G, O-Q). (A-Q) Scale bar, 200μm.

We then examined the role of Notch signaling in *PAPC* regulation in chicken embryo explants using the γ-secretase inhibitor DAPT. Treatment resulted in the complete loss of *PAPC* expression stripes in the anterior PSM (Fig. 2A, C, D; n=9). However, in the DAPT-treated embryos, *PAPC* maintained a distinct cycling expression domain in the posterior PSM (Fig. 2C, D). This suggests that *PAPC* dynamic expression in the posterior PSM does not depend on Notch signaling. Overexpression of the chicken *Mesp2* homolog (*MESO2*) by electroporation, resulted in a strong ectopic expression of *PAPC* throughout the paraxial mesoderm (Fig. 2G; n=7) compared to control embryos (Fig. 2E; n=8). Together, these data confirm that in the anterior PSM of the chicken embryo, *PAPC* becomes regulated by *Mesp2/MESO2* downstream of Notch signaling as reported in frog and mouse (Kim et al., 2000; Rhee et al., 2003) ( Nomura-Kitabayashi et al., 2002).

To assess if the regulation of PAPC expression is conserved in amniotes, we reinvestigated *PAPC/Pcdh8* mRNA and protein expression in mouse embryos (Makarenkova et al., 2005; Rhee et al., 2003; Yamamoto et al., 2000). We generated a polyclonal antibody against the mouse PAPC protein and observed an antigen distribution similar to that observed in the chicken paraxial mesoderm. Mouse PAPC is expressed in dynamic stripes in the anterior PSM and becomes localized in the rostral compartment of the forming somite S-I, posteriorly to the forming boundary (Fig. 1H). While *PAPC* mRNA expression is much fainter in the posterior PSM than in the anterior PSM, analysis of stage-matched mouse embryos showed distinct patterns of *PAPC* mRNA expression in the posterior PSM (arrowheads, Fig. 2H-J), consistent with the idea that *PAPC* expression is also regulated by the Segmentation Clock in mouse.

We then examined *PAPC* expression in several mouse mutants for key signaling pathways involved in the Segmentation Clock control and PSM maturation. In *RBP-jK* mutant mice that are defective for Notch signaling (Oka et al., 1995), *PAPC* expression was strongly reduced, but one or two diffuse expression stripes were nevertheless observed in a region completely lacking somites in the posterior paraxial mesoderm (arrowheads, Fig. 2K, L; n= 17). These data suggest that, as in chicken embryos, the periodic activation of mouse *PAPC* in the posterior PSM is independent of Notch signaling. The Vestigial tail (*Vt*) mouse mutant is a hypomorphic mutant of *Wnt3a*, which exhibits a loss of Wnt signaling in the tail bud at E10.5 dpc (Aulehla et al., 2003; Greco et al., 1996). In this mutant, *PAPC* expression was strongly down-regulated, and the expression stripes were often fused and mispatterned compared to control (arrowheads, Fig. 2M-O; n=5). Retinoic acid signaling has also been shown to contribute to paraxial mesoderm maturation by antagonizing the Wnt/FGF posterior gradient (Diez del Corral et al., 2003; Moreno and Kintner, 2004; Niederreither and Dolle, 2008). In the *Raldh2* null mutant mice which are defective for retinoic acid production, *PAPC* expression was restricted to the anterior-most PSM and formed narrow asymmetrical stripes (Fig. 2P, Q; n=5). As *Raldh2* mutants exhibit a FGF signaling gain-of-function phenotype in the PSM (Vermot et al., 2005), the lack of posterior expression of *PAPC* in these mutants is consistent with the FGF-dependent repression observed in chicken embryos. Thus our data identifies *PAPC/Pcdh8* as a novel type of cyclic gene exhibiting an unusual periodic repression in the PSM downstream of FGF signaling in the mouse and chicken embryo.

### Complementary distribution of PAPC and CDH2 along the PSM

In the anterior PSM, although CDH2 is present throughout the tissue, its protein exhibits a striking segmental pattern complementary to PAPC (Fig.3A-B). In the chicken embryo, epithelialization of the posterior somitic wall begins before the fissure forms (Duband et al., 1987; Nakaya et al., 2004). Thus, in the forming somite (S0), the epithelialization process is more advanced in the caudal compartment than in the rostral one, as evidenced by increased co-localization of CDH2 and F-Actin (Fig. 3B, F). The rostral compartment of S0 acquires an epithelial polarity and becomes recruited to the rosette soon after formation of the posterior fissure. This transition correlates with the down-regulation of PAPC in the rostral compartment of the newly formed somite, and with the accumulation of CDH2 and F-Actin at the apical membrane of anterior somitic cells (Figure 3D, F). At the forming boundary level (S-I/S0), PAPC is excluded from the posterior epithelializing domain of S0 (Fig. 1I, 3C-E). In the posterior wall of S0, CDH2 was found to be essentially located at the cell membrane of epithelial cells (Fig. 3D). In contrast, in the rostral part of S-I, CDH2 and PAPC are also found intracellularly. Using the anti-PAPC polyclonal antibody and a secondary antibody labeled with gold particles, we analyzed PAPC distribution by electron microscopy in the anterior compartment of S0 and detected PAPC primarily at cell-cell contacts and sites of membrane trafficking, including clathrin pits and endocytosis vesicles (Fig.S1).

**Fig. 3.**
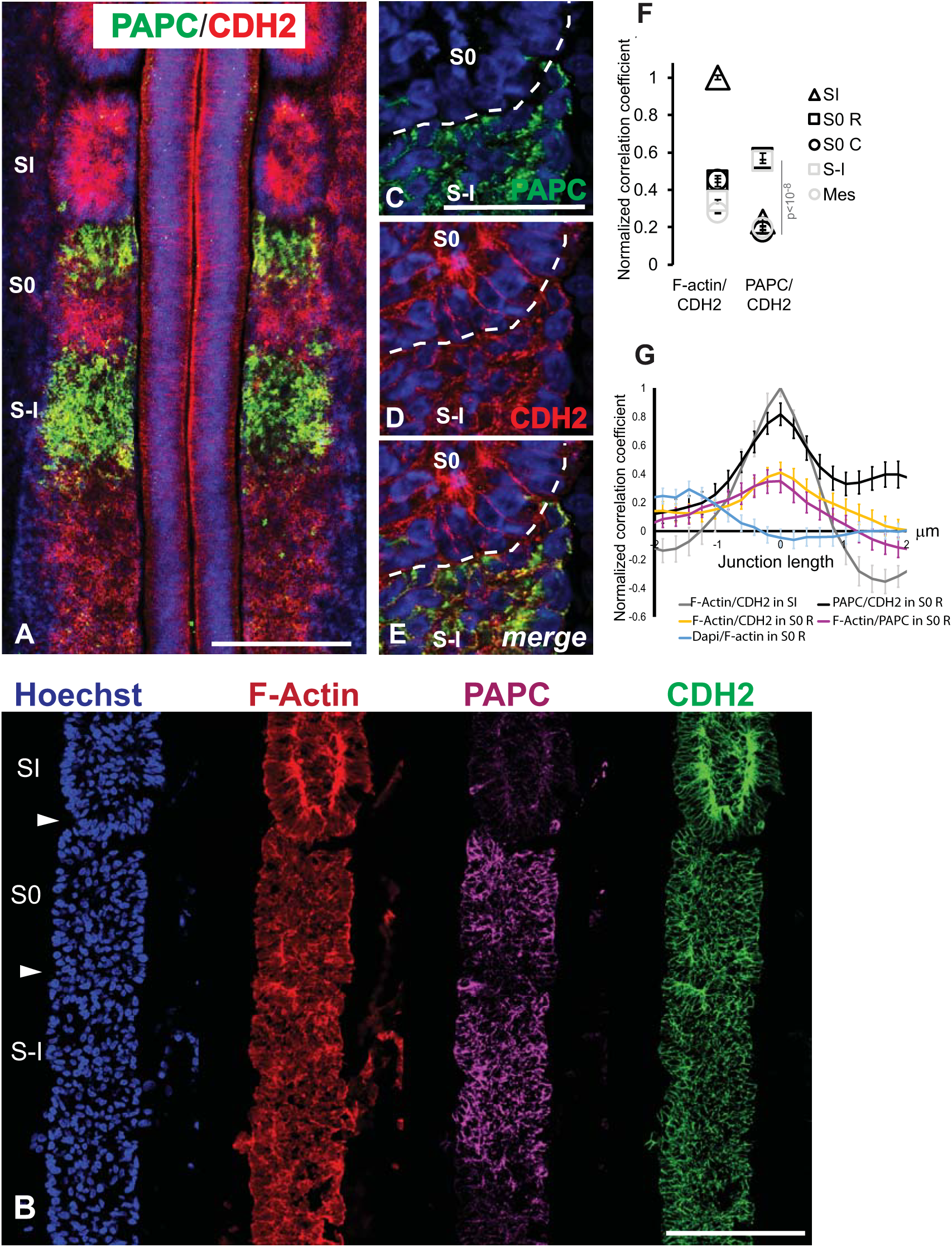
Complementary distribution of PAPC and CDH2 in the forming somite. **(A)** Comparison of PAPC and CDH2 distribution during somite formation in the anterior PSM. Whole-mount immunohistochemistry with PAPC antibody (green) and CDH2 antibody (red). Dorsal view, anterior to the top. Nuclei are labeled in blue. Scale bar, 200μm. **(B)** Confocal image of immunostainings of parasagittal sections of the PSM of a stage 15 somite embryo showing the localization of nuclei (Hoechst, blue), F-actin (Phalloidin, red), PAPC (magenta) and CDH2 (green). Scale bar, 100μm. **(C-E)** Parasagittal cryosection showing a higher magnification of the region of the interface between S0 and S-I immunostained for PAPC (C, green) and CDH2 (D, red) and merge (E). The dashed white line marks the position of the forming boundary. Nuclei are labeled in blue. Scale bar, 50μm. **(F)** PAPC, CDH2 and F-Actin colocalization analysis in the anterior PSM.
Normalized correlation coefficients were calculated based on signal intensity profiles for each staining within each PSM subdomains (n=3 embryos). For each 2 by 2 comparison, a Pearson’s coefficient was calculated with 1 corresponding to a total positive correlation. Each sub-region of the PSM has been divided in 5 area of 19 um^2^. S-I/0/I: somite -I/0/I; R: rostral half; C: caudal half; mes.: posterior PSM mesenchyme. p: p-value. **(G)** PAPC, CDH2 and F-Actin colocalization analysis at cellular junctions. Normalized correlation coefficients were calculated based on signal intensity profiles along cell-cell junctions (over 3μm in length) in various PSM subdomains (S0/I: somite 0/I; R: rostral.). For each 2 by 2 comparison, a Pearson’s coefficient was calculated with 1 corresponding to a total positive correlation, A peak at 0 micron means that both signals are co-localized. n= 20 junctions per domain.

In order to evaluate CDH2 and PAPC colocalization, we performed a cross-correlation study of the intensity profile of PAPC and CDH2, which was compared to a similar analysis between CDH2 and F-Actin (which colocalize in the apical domain of epithelial cells). The analysis was performed in the anterior PSM (Fig.3B, F, Fig.S2). As expected, the strongest correlation coefficient obtained was for CDH2 and F-Actin in the newly formed somite (SI) where CDH2 complexes are stabilized by the network of F-Actin to maintain the epithelial structure of the somite (Fig. 3F). Interestingly, we observed a graded decrease of the correlation coefficient of CDH2 and F-Actin from high in somite to low in the PSM consistent with the progressive nature of the epithelialization process along the PSM (Fig. 3F, Fig.S2). The correlation between PAPC and CDH2 was maximal in the rostral part of the Somite 0 (S0 R) and in the Somite -I (S-I) where PAPC is strongly expressed indicating that CDH2 and PAPC largely co-localize in these regions of the PSM (Fig. 3B, F).

We also performed cross-correlation measurements of the intensity profile of PAPC and CDH2 or of CDH2 and F-Actin at the interface between cells in the rostral part of S0 (Fig. 3G, Fig. S2). As in the cross-correlation analysis described above, the values were compared to that obtained for F-Actin and CDH2 colocalization in the somite. Interestingly, the correlation was high (corr. coef = 0.8) with a peak of colocalization found at 0 micron indicating that PAPC and CDH2 are strongly colocalizing in the rostral part of S0, at a 200nm resolution (Fig. 3G). These data suggest that PAPC and CDH2 are also located in close proximity at the cell membrane in the rostral compartment of S0. Attempts to co-immunoprecipitate PAPC with CDH2 from chicken PSM extracts were negative, suggesting that the PAPC/CDH2 do not directly interact (data not shown).

### PAPC regulates somite boundary formation and CDH2 distribution at the cell membrane

The striking complementary patterns of PAPC and CDH2 expression during somitogenesis prompted us to examine whether PAPC interferes with CDH2 function during somitogenesis. We used *in ovo* electroporation of the primitive streak paraxial mesoderm precursors to overexpress PAPC constructs in the PSM (Dubrulle et al., 2001). Embryos overexpressing the long PAPC isoform (PAPC-L) were not different from control embryos overexpressing the empty vector (data not shown). In contrast, when we overexpressed the short PAPC isoform (PAPC-S) and a membrane-bound GFP reporter, electroporated cells formed clumps of cells, which interfered with proper somite morphogenesis, forming mesenchymal bridges that blocked boundary formation (Fig. 4A-B, n= 25, control n=15). Intersomitic fissures were often lacking in PAPC-S overexpressing embryos and somites were not properly separated (Fig. 4B). Also electroporated anterior PSM and newly formed somite cells exhibited a rounder, more mesenchymal morphology, losing the apical accumulation of CDH2 (Fig. 4D-I). We also observed differential sorting of the electroporated cells which were found preferentially in the rostral mesenchymal compartment of newly formed somites (Fig. 4J, K; n= 25). In clumps of PAPC-S overexpressing cells, CDH2 expression was only detectable at a low level at the cell membrane compared to neighboring cells and to control electroporated cells (compare Fig. 4D-F and G-I). Hence, overexpression of PAPC-S in anterior PSM cells reduces CDH2 distribution at the membrane of the expressing cells. CDH2 plays an important role in the maintenance of the epithelial structure in anterior PSM (Duband et al., 1987; Horikawa et al., 1999; Linask et al., 1998; Radice et al., 1997) and in boundary formation (McMillen et al., 2016). Thus, the reduction of CDH2 at the membrane of cells overexpressing PAPC could explain their loss of epithelial polarity, their acquisition of a mesenchymal fate and their segregation in the rostral compartment of the forming somite.

**Fig. 4.**
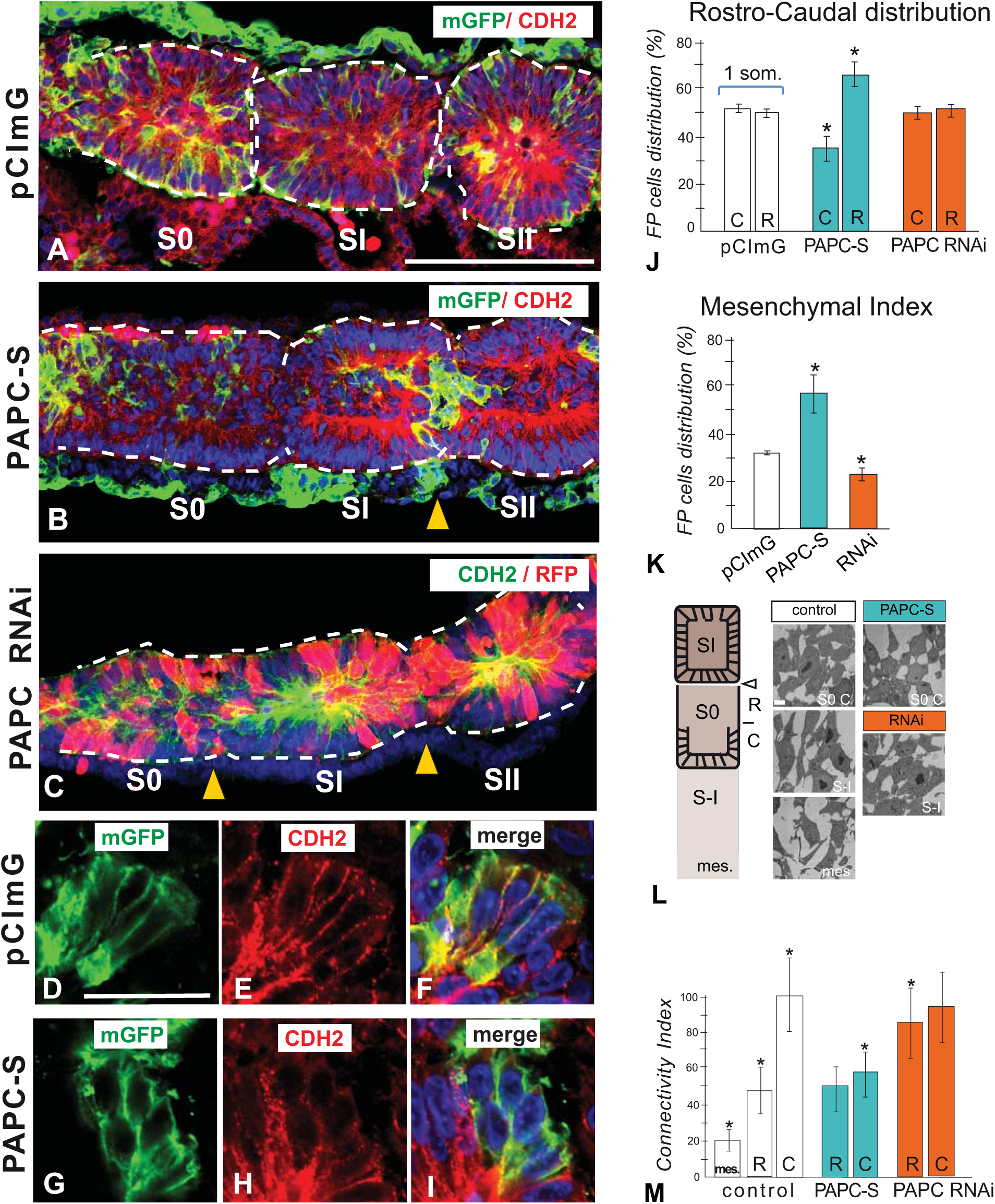
*PAPC* overexpression disrupts somite boundary formation and CDH2 localization. **(A-I)** CDH2 immunostaining of parasagittal sections of the PSM of E2 chicken embryos electroporated with pCImG (control, A), PAPC-S (B), and PAPC-RNAi (C). Nuclei are shown in blue. Electroporated cells coexpress a membrane-bound GFP (green, A, B, D, F, G, I) or RFP (red, C). Somite limits are highlighted by dashed white lines. Arrowheads denote segmentation defects. (A-C) Scale bar, 100μm. (D-I) Higher magnification of the newly formed somite SI of E2 chicken embryos electroporated with the pCImG (D, I) and pCImG-PAPC-S constructs showing immunostainings of parasagittal cryosections labeled with an anti-GFP (green, D, G), and anti-CDH2 (red, E, H) and merged panels (F, I). (D-I) Scale bar, 25μm. **(J)** Quantification of the electroporated cell distribution along the rostral [R] and caudal [C] somite compartments in chicken embryos electroporated with pCImG, PAPC-S, and PAPC-RNAi constructs, respectively. **(K)** Mesenchymal index as defined by the distribution of electroporated cells in the mesenchymal and epithelial fraction of the newly formed somites in embryos electroporated with pCImG, PAPC-S, and PAPC-RNAi constructs, respectively. **(L)** Electron Microscopy (EM) analysis of cellular organization in the PSM of chicken embryos electroporated with pCImG, PAPC-S and PAPC-RNAi constructs. (left) PSM diagram indicating the sites of analysis, namely posterior PSM (mesenchyme (mes.)), the rostral compartment of S-I [R] and caudal compartment of S0 [C] at the level of the forming boundary. (right) representative EM pictures of the cellular organization of control and treated domains. Scale bar, 2μm. **(M)** Cell-cell connectivity index in control embryos, PAPC-S and PAPC-RNAi electroporated embryos, respectively. (J,K,M) Mean +/− s.e.m. * p<0.05, refer to Materials and Methods for quantification method

We then examined the effect of PAPC knock-down in anterior PSM cells. Embryos electroporated with PAPC-RNAi exhibited strong down-regulation of *PAPC* mRNA expression (Fig. S3), associated with a loss of PAPC protein in expressing cells compared to controls (Fig. 1B, Fig. S3). Notably, the PAPC-RNAi-expressing cells were located preferentially in the epithelial fraction of the somite, showing an opposite behavior to cells expressing PAPC-S (Fig. 4C, K, n=25). Cells in which PAPC was knocked-down were preferentially located in the epithelial layer and acquired a spindle-shape morphology, accumulating CDH2 and actin cytoskeleton at their membrane (Fig. 4C). These cells formed epithelial connections between somites, interfering with proper segmentation and somite morphogenesis. No cell distribution bias along the rostro-caudal compartments of somites was observed, possibly due to the increase in cell adhesivity (Fig. 4J). These data support the idea that PAPC activity regulates the membrane distribution of CDH2, promoting a more mesenchymal state.

We further analyzed the effect of overexpressing PAPC-S and PAPC-RNAi constructs on the cytoarchitecture of paraxial mesoderm cells (Fig. 4L). By electron microscopy, a clear transition was observed on both sides of the future somitic boundary, between the mesenchymal nature of S-I and the epithelial organization of the posterior wall of S0 (Fig. 4L). The progressive epithelialization of the anterior PSM was characterized by a large increase in cell-cell connectivity, namely an increase in the number of cell-cell contacts and their length (Fig. 4L, M). Overexpression of PAPC-S resulted in a significant decrease in cell-cell connectivity in the caudal compartment of S0 compared to control (Fig. 4L, M; n=4). Conversely, expression of PAPC-RNAi constructs in the anterior compartment of S-I resulted in an increase of cell connectivity (Fig 4L, M; n=4). This suggests that the expression level of PAPC in PSM cells is inversely correlated to their epithelialization status.

To determine whether PAPC antagonizes CDH2 function, we attempted to rescue the PAPC-S overexpression phenotype by co-electroporating a CDH2-expressing construct. However, direct expression of CDH2 in the epiblast leads to cell-sorting defects during gastrulation, thus preventing the analysis of the PSM phenotypes (data not shown). To circumvent this, we used an inducible system to restrict overexpression of CDH2 once the cells entered the PSM. The phenotype of embryos overexpressing only CDH2 in the anterior PSM resembled that of PAPC loss-of-function with overexpressing cells clustering and integrating the epithelial compartment of the anterior PSM (Fig.5A-H, n=25). In these embryos, intersomitic fissures were often absent as cells remained connected by epithelial bridges (Fig. 5E). Co-electroporation of the inducible CDH2 construct together with PAPC-S led to a partial rescue of somite morphogenesis (Fig.5I, n=32). In these embryos, somite morphology was partially restored, and co-electroporated cells contributed both to the somite epithelial ring and to the mesenchymal somitocoele compartment (Fig.5I-M). These experiments suggest that PAPC can antagonize the epithelialization-promoting function of CDH2 in the anterior PSM.

**Fig. 5.**
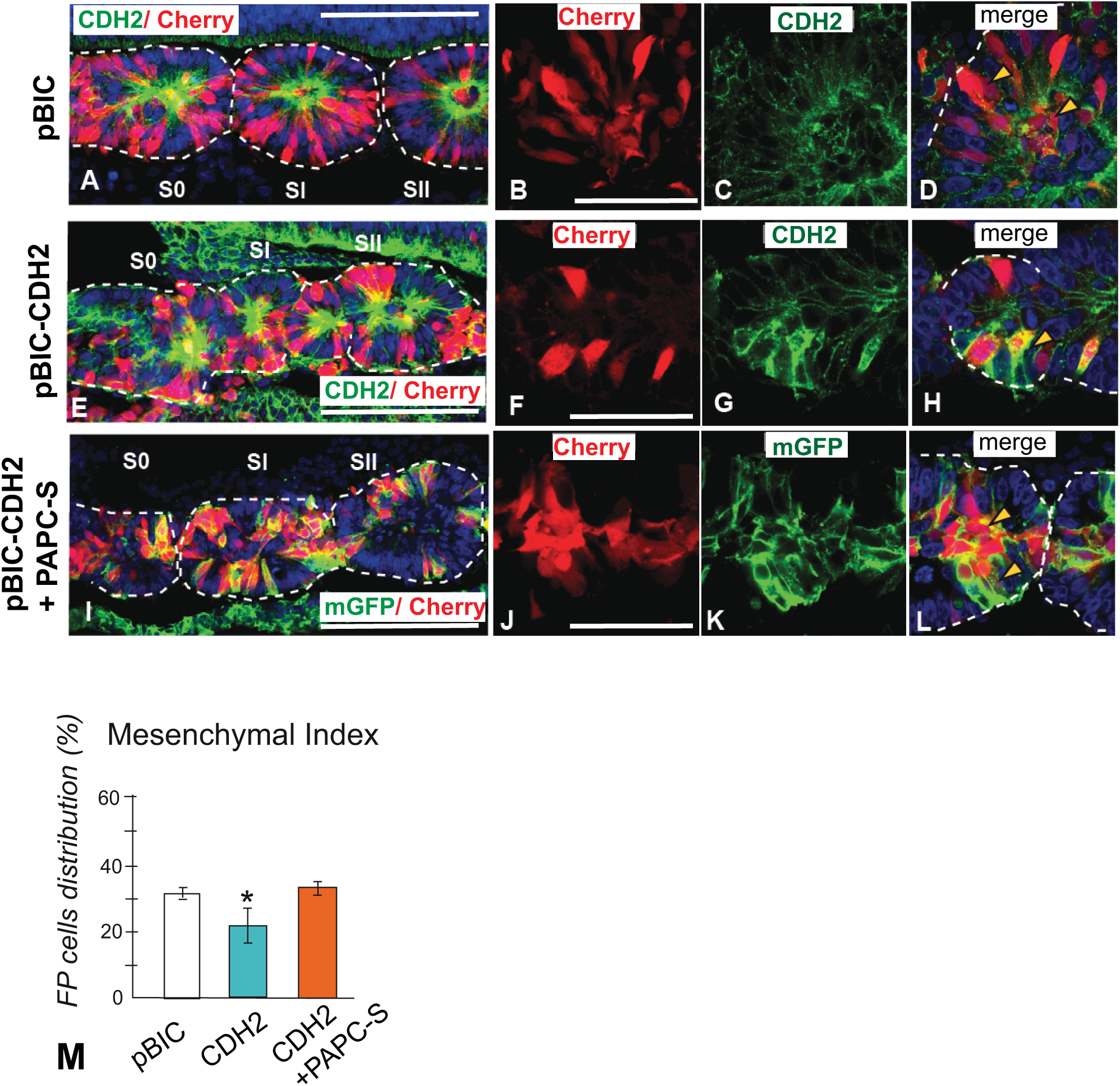
PAPC negatively regulates CDH2 during somite morphogenesis. **(A-L)** Immunostainings of parasagittal cryosections of the PSM of E2 chicken embryos electroporated with pBIC (control, A-D), pBIC-CDH2 (E-H), or coelectroporated with pBIC-CDH2 and pCImG-PAPC-S (I-L). Electroporated cells with pBIC vectors coexpress mCherry (red), while pCImG vector coexpress membrane-bound GFP (green, I-L). Nuclei in A, E, I and D, H, L are shown in blue. Somite individualization is highlighted by dashed white lines. **(A,E,I)** Low magnification images. Scale bar, 100μm. **(B-D, F-H, J-L)** Higher magnification of parasagittal cryosections of the forming somite region of E2 chicken embryos electroporated with pBIC (B-D), pBIC-CDH2 (F-H), and pBIC-CDH2 + pCImG PAPC-S (J-L) showing immunostainings labeled with an anti-Cherry (Red, B, F, J), an anti-CDH2 (green, C, G), an anti-mGFP (green, K) and merged panels (D, H, L). Scale bar, 25μm. **(M)** Mesenchymal index as defined by the distribution of electroporated cells in the mesenchymal and epithelial fraction of the newly formed somites electroporated with pBIC, pBIC-CDH2, and coelectroporated with pBIC-CDH2 + pCImG PAPC-S, respectively. Mean +/− s.e.m. * p<0.05.

### PAPC regulates endocytosis in the anterior compartment of the forming somite

Our data suggest that interfering with PAPC function can alter the epithelial state of PSM cells and the level of CDH2 at the cell membrane. In the rat nervous system, PAPC/Arcadlin was shown to regulate CDH2 function by controlling its endocytosis (Yasuda et al., 2007). This led us to ask whether PAPC regulates CDH2 distribution by regulating clathrin-dependent endocytosis in the paraxial mesoderm. Interestingly, the rostral compartment of S-I, which strongly expresses PAPC, shows higher levels of punctate clathrin staining, suggesting more active endocytosis compared to the newly formed somite S0 (Fig. 6A-F). In the rostral compartment of S-I, PAPC and CDH2 distribution largely overlaps with clathrin (Fig.6G-L).

**Fig.6.**
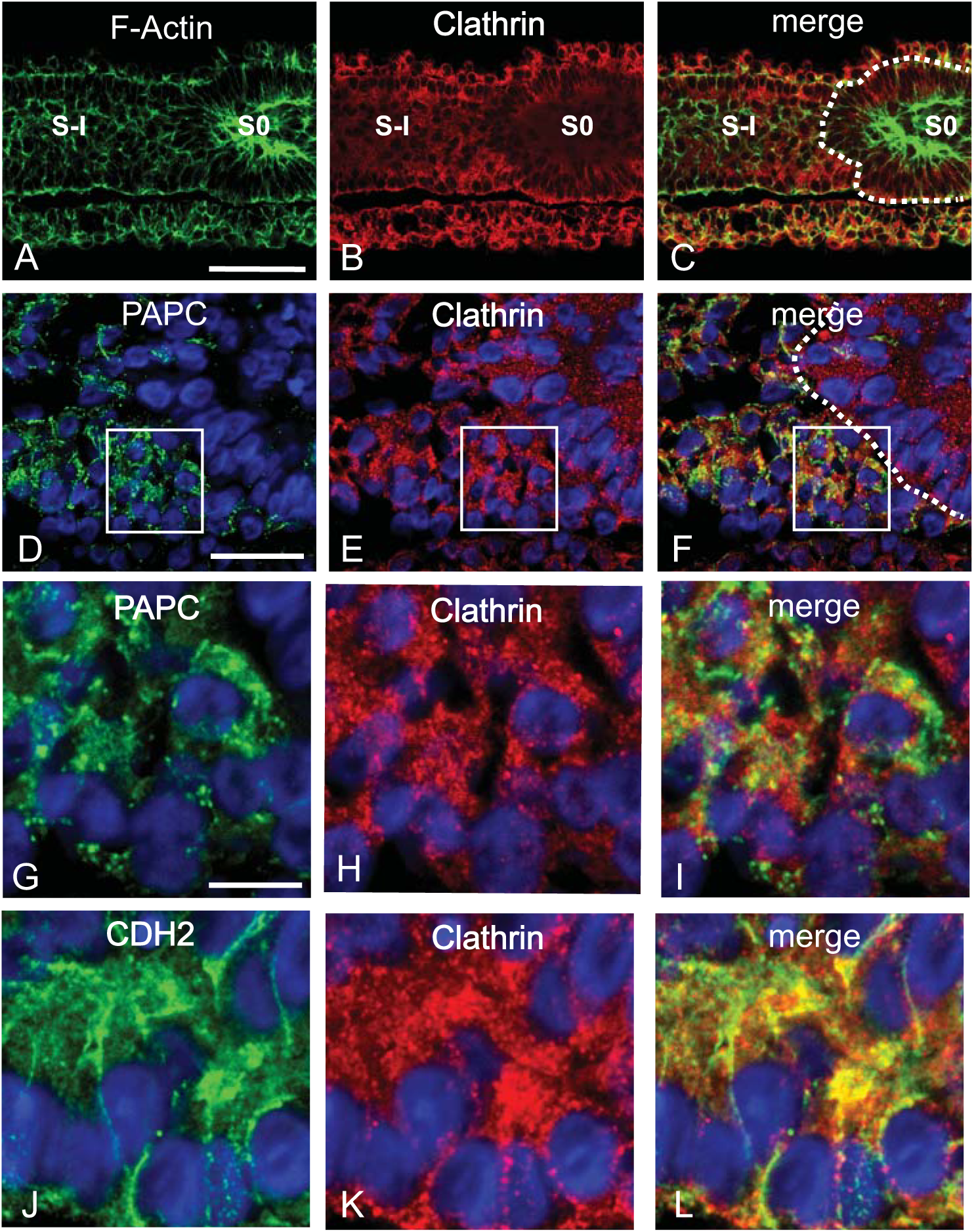
PAPC, CDH2 and clathrin colocalize in the anterior PSM (A-C) **(A-C)** Parasagittal sections of the forming posterior boundary of S0 labeled with phalloidin (A, F-actin, green), and an antibody against clathrin (B, red). (C) Overlay of the panels A and B. Scale bar, 50μm. **(D-L)** Higher resolution images showing the comparison of Clathrin, PAPC (D-I) and CDH2 (J-L) proteins distribution during somite formation. Panels (G-I) are detail of panels (D-F) (white boxes). Proteins are detected by immunofluorescence and nuclei are counterstained (blue). Parasagittal sections, anterior to the right. Scale bar, 20μm (D-F), 10μm (G-L).

In order to test if PAPC can activate the clathrin-dependent endocytosis pathway, we compared the uptake of fluorescent dextran in embryos electroporated with an empty GFP expressing construct (empty pCImG vector) or with the same vector containing the PAPC-S coding sequence (Fig. 7A, B; Fig. S4). We quantified the intensity of PAPC and of dextran fluorescence in GFP+ cells (carrying the vector) and in GFP-cells (Fig. 7A, B; Fig.S4). The ratio of fluorescence intensity between GFP positive and GFP negative cells was found increased for PAPC in cells overexpressing the PAPC-S construct as expected. These cells also exhibited an increase in dextran fluorescence (Fig 7A,B; Fig. S4). Treating the electroporated embryos with Pitstop2 (an inhibitor of clathrin-mediated endocytosis), prior to the addition of dextran, significantly decreased the uptake of Dextran in PAPC-S overexpressing cells (Fig. 7A-B, Fig.S4). These experiments therefore suggest that PAPC can increase the rate of clathrin-mediated endocytosis.

**Fig.7.**
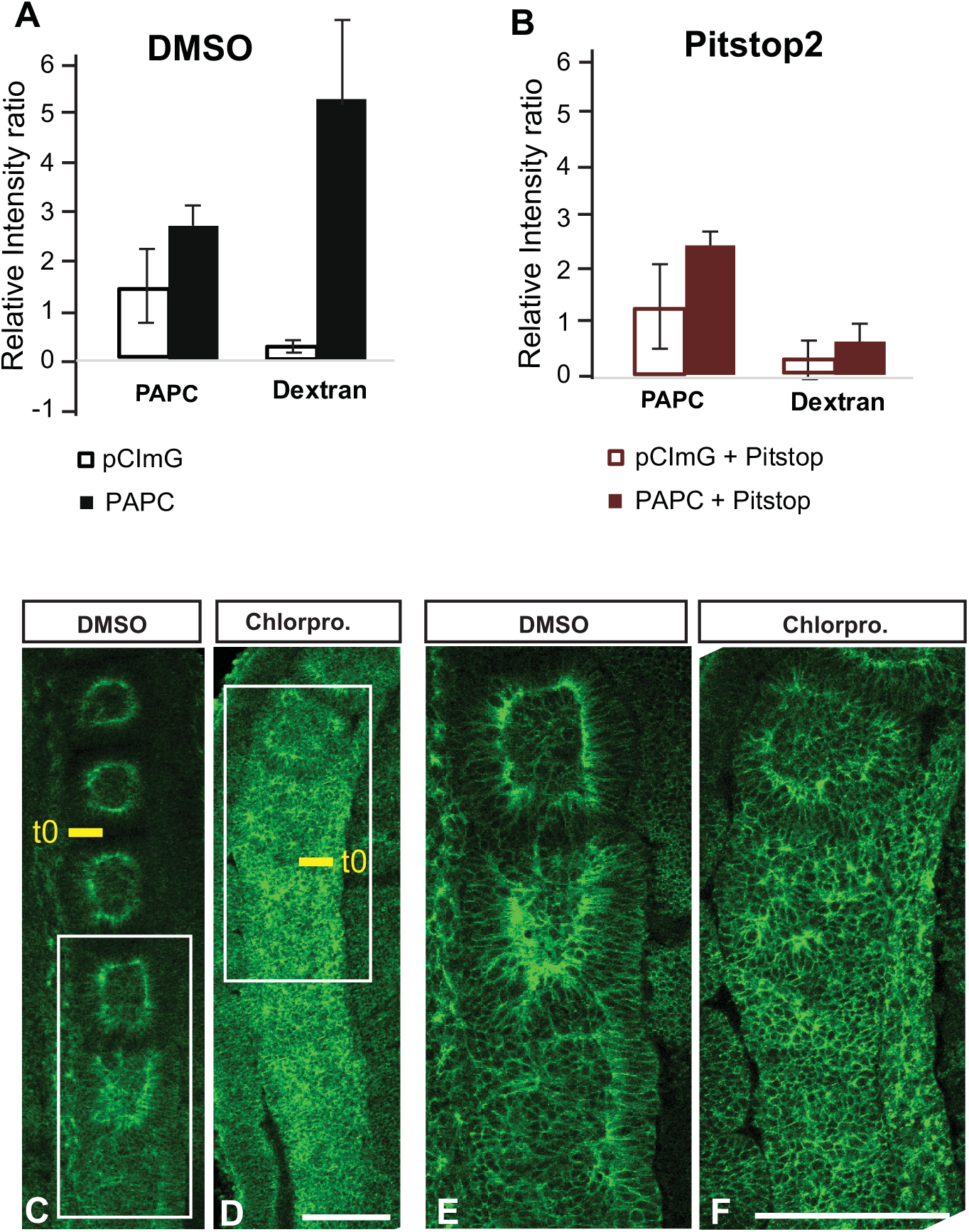
PAPC promote anterior PSM cells endocytic activity. (A-B) Quantification of dextran uptake as a measure of endocytosis level. Fluorescence intensity for PAPC and Dextran was measured in embryos electroporated with pCImG (empty histograms) or PAPC-S (full histograms) and subsequently treated with DMSO (control) or Pitstop2 (red). The fluorescence intensity ratio of the GFP+ over GFP-cells are shown. Number of embryos analyzed per conditions: pCImG/DMSO n=2; pCImG/Pitsop2 n=4; PAPC-S/DMSO n=7; PAPC-S/Pitstop2 n=6. Mean +/− s.d. **(C-F)** CDH2 distribution in chicken PSM explants cultured 4 hours in the presence of DMSO (0.2%) (C, E) and Chlorpromazine (50μM) (D, F). Panels E and F correspond to the boxed area shown in C and D, respectively. t0: last formed boundary at treatment start. Dorsal view, anterior to the top. Scale bar, 100μm.

To further test the role of endocytosis in somite boundary formation, we treated cultured embryo explants with the endocytosis inhibitor Chlorpromazine for 3-5 hours. While control explants formed several somites (Fig. 7C, E), treated explants did not form any new somites (Fig. 7D, F; n=7). PSM cells of treated explants failed to adopt an elongated polarized epithelial morphology, as evidenced by the failure of ZO-1 accumulation and of tight junctions formation (Fig. S5). Moreover, treated explants showed little signs of epithelial polarization, displaying a higher level of membrane CDH2 (Fig.7F). These data support the idea that PAPC is involved in active CDH2 endocytosis, and that this process is critical for the formation of the somitic fissure (Fig.8).

**Fig.8.**
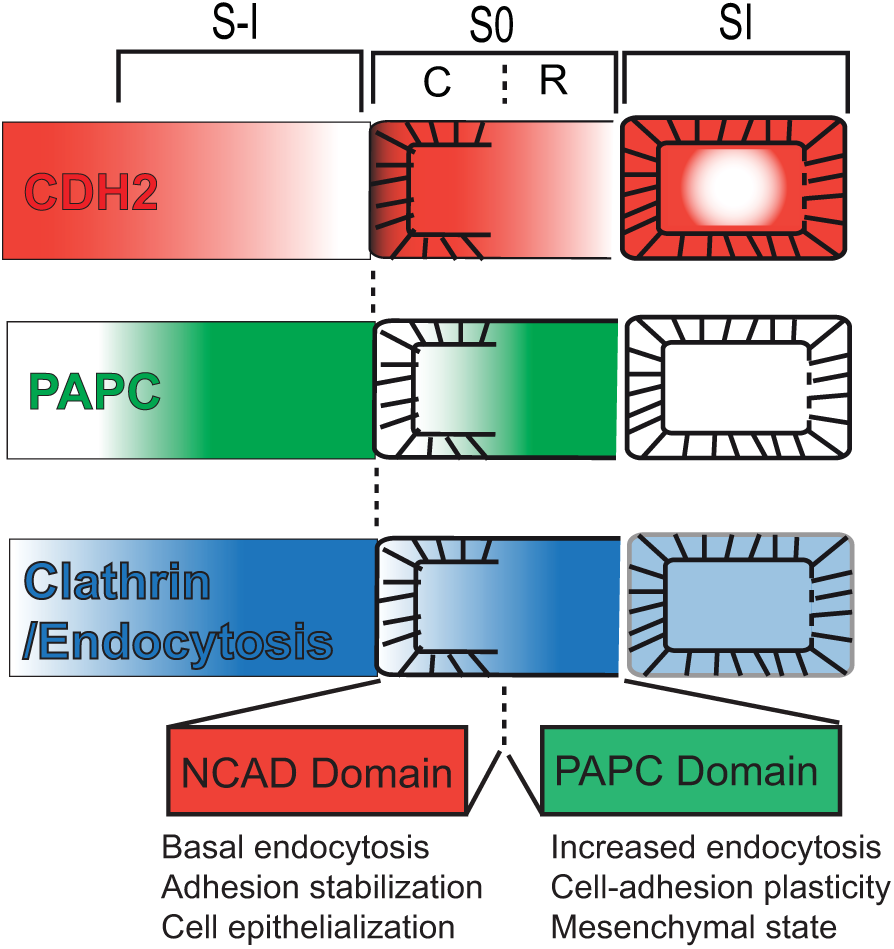
Model for the role of PAPC in somite segmentation. Segmental expression of PAPC protein (green) is superimposed on the CDH2 adhesion field (red) leading to enhanced endocytosis (blue) clearing locally CDH2 from the cell surface and generating a de-adhesion interface, which allow the formation of a new somitic boundary.

## DISCUSSION

How the rhythmic signal of the Segmentation Clock translates into the activation of a periodic morphogenetic program, ultimately leading to the formation of the epithelial somites is not well understood. Here, we provide evidence for a direct coupling between the Segmentation Clock oscillator and somite morphogenesis via the periodic regulation of the protocadherin PAPC, which increases the clathrin mediated endocytosis dynamics of CDH2 to control the formation of the posterior somitic fissure.

*PAPC* expression has been described in the anterior PSM of fish, frog and mouse embryos where it is expressed as dynamic stripes (Kim et al., 2000; Rhee et al., 2003; Yamamoto et al., 1998). We show that *PAPC* expression in the posterior PSM shows periodic waves of gene expression with similar kinetics to the *LFNG* waves associated to the Segmentation Clock (Forsberg et al., 1998; McGrew et al., 1998). Blocking Notch signaling pharmacologically in the chicken embryo or genetically in the mouse does not interfere with *PAPC* dynamic expression in the posterior PSM indicating that this periodic regulation is independent of Notch. Overexpression of the bHlH transcription factor *MESPO* (the homologue of Mesogeninl a key target of the Wnt pathway in the PSM (Wittler et al., 2007)) is sufficient to drive ectopic *PAPC* expression in the mesoderm in the chicken embryo. Together with the *PAPC* downregulation observed in the mouse *Wnt3a* mutant (*Vt*), this suggests that *PAPC* acts downstream of the Wnt signaling pathway in the posterior PSM. We show that in the posterior PSM, *PAPC* expression is negatively regulated by FGF signaling and exhibits an expression gradient opposite to the FGF gradient. Since FGF signaling has been shown to be periodically activated in the posterior PSM in mouse and chicken embryos (Dale et al., 2006; Dequeant et al., 2006; Krol et al., 2011; Niwa et al., 2007), this suggests that *PAPC* is periodically repressed by FGF signaling. Thus, our data indicate that *PAPC* is a novel cyclic gene, periodically repressed by FGF signaling in the posterior PSM of chicken embryos.

We show that Notch inhibition prevents the formation of *PAPC* stripes in the anterior PSM. In mouse, *PAPC* expression is lost in the *Mesp2*-/- mutant (Nomura-Kitabayashi et al., 2002; Rhee et al., 2003) whereas, in frog and chicken embryos, overexpression of the *Mesp2* homologues, leads to ectopic activation of *PAPC* (Kim et al., 2000). Furthermore, *PAPC* expression in the anterior PSM tightly overlaps with the *Mesp2* stripes. Together, these data show that *PAPC* is a conserved target of the Mesp2 transcription factor and acts downstream of Notch signaling in the anterior PSM in vertebrates.

Our data suggest that in the chicken embryo, PAPC prevents epithelialization of cells in the rostral somite compartment by controlling CDH2 endocytosis, thus negatively regulating its function. This results in a sawtooth pattern of CDH2 resembling that recently described in zebrafish (McMillen et al., 2016). The action of PAPC on CDH2 endocytosis leads to the establishment of an interface between cells expressing high levels of CDH2 (posterior S0) and lower CDH2 levels (anterior S-I) at their cell membrane at the forming somite border. In zebrafish, CDH2 inhibits integrin alpha5 activation in adjacent PSM and such an interface is required to allow the activation of integrin alpha5 by the EPHA4-EphrinB2 system (Julich et al., 2009;Koshida et al., 2005; McMillen et al., 2016). Integrin activation at this interface results in the local assembly of fibronectin fibrils in the forming intersomitic fissure. Our results provide a possible mechanism for the establishment of this important interface between the mesenchymal domain of the rostral part of S-I and the epithelial domain of the caudal part of S0. Such an interface could promote the de-adhesion behavior involved in somite boundary formation and the subsequent matrix-filled fissure formation (Fig. 8). No somitic defects have however been reported in mice mutant for *EphA4*, *EphrinB2*, and *PAPC* (Adams et al., 1999 ; Dottori et al., 1998; Yamamoto et al., 2000), supporting some level of functional redundancy among these different pathways involved in somite boundary formation.

Several studies suggest that PAPC indirectly controls cell adhesion by negatively regulating the function of classical cadherins (Chen and Gumbiner, 2006; Chen et al., 2009; Yasuda et al., 2007). CDH2 is a major component of adherens junctions, which has been implicated in somite morphogenesis (Duband et al., 1987; Horikawa et al., 1999; Radice et al., 1997; Linask et al., 1998). Our study demonstrates that PAPC and CDH2 colocalize at cell-cell junctions and also in trafficking vesicles in the anterior compartment of the forming somites. Remarkably, where the two proteins are coexpressed, CDH2 appears abundant in the cytoplasm of the cells and at loose cell-cell connections; whereas, in domains lacking PAPC expression such as the caudal domain of the forming somite, CDH2 essentially localizes at the cell membrane. Moreover, cells overexpressing PAPC in the anterior PSM lose their epithelial structure and downregulate CDH2 expression at the cell membrane. PAPC overexpression resulted in a striking cell-sorting phenotype and a disruption of normal boundary formation, consistent with a modulation of the CDH2-dependent adhesion of overexpressing cells in the anterior PSM. CDH2 overexpression also results in expressing cells to adopt an epithelial fate. This effect can be partly rescued by overexpressing PAPC together with CDH2, supporting the idea that PAPC antagonizes CDH2 function in the anterior PSM. Endocytosis has been shown to modulate the adhesive properties of cadherins (Troyanovsky et al., 2006). Interestingly, in the nervous system, PAPC (Arcadlin) has been shown to directly trigger CDH2 endocytosis through a p38 MAPK activation, leading to dendrite remodelling (Yasuda et al., 2007). Our results also show that clathrin mediated endocytosis is active in the anterior PSM and becomes overactivated in PAPC electroporated cells. Together, these observations support a model in which PAPC antagonizes CDH2 function in the rostral part of the forming somite by promoting its endocytosis. As a result, cells of the rostral compartment of the somite S-I remain mesenchymal; whereas cells of the caudal compartment of the somite S0 form an epithelial posterior wall. This interface will form the posterior somitic fissure (Fig. 8). While the PSM exhibits an overall uniformly graded distribution of CDH2 in the anterior PSM, our work suggest that the periodic regulation of CDH2 trafficking mediated by PAPC downstream of the Segmentation Clock triggers local de-adhesion, creating the interface forming the somitic fissure (Fig. 8). Thus, PAPC functions as a morphogenetic translator of the input of the Clock and Wavefront system, leading to periodic somite boundary formation.

## Acknowledgements

We thank Jennifer Pace, Leif Kennedy, Merry McLaird and Barbara Brede for excellent technical help, members of the Pourquié laboratory for helpful discussions. We thank Alexis Hubaud for critical reading of the manuscript. We are grateful to the Stowers Institute Core Centers, to Fengli Guo (Stowers) and Yannick Schwab (IGBMC) for Electron Microscopy. We thank Joanne Chatfield for manuscript editing and Silvia Esteban for artwork. We thank T. Honjo, P. Chambon, and S.A. Camper for providing the *RBPjK*, *Raldh2*, *Vt mice*, respectively. This work was supported by Stowers Institute for Medical Research and Howard Hughes Medical Institute. This work has been partly supported by a NIH grant R01 HD043158 to O.P. and by an advanced grant from the European Research Council to O.P.

## Author contributions

J.C. designed and performed most of the experiments, analyzed data. C.G. performed and analyzed the immunolocalization and endocytosis assays. O.P. supervised the overall project. J.C., C.G, O.P. performed the final data analysis and wrote the manuscript.

## Competing financial interests

The authors declare no competing financial interests that might be perceived to influence the work reported in this article.

## SUPPLEMENTARY MATERIAL

### FIGURE LEGENDS

**Figure S1**

Subcellular localization of PAPC in anterior PSM cells by IEM. PAPC can be found (gold beads, black dots) specifically at cell-cell junctions and sites of membrane trafficking including clathrin coated pits and endocytosis vesicles. Scale bar, 200nm.

**Figure S2**

**(A)** (Left) Diagram of the PSM subdomains. S-I/0/I:somite -I/0/I; R: rostral half; C: caudal half; mes.: posterior PSM mesenchyme. (Right) Representative confocal sections for each subdomains as shown on the diagram (SI, top row; mes., bottom row), after co-staining for F-Actin (Phalloidin, red), PAPC (purple), CDH2 (green). Note that PAPC is detected only in PSM subdomains S-I and S0. Each image shows a representative image of the 19μm^2^ areas used for the quantification shown in Fig. 3F. Arrows: F-Actin and CDH2 co-localization; Arrowheads: PAPC and CDH2 co-localization. Scale bar, 4μm.

**(B, C)** Cell-cell junction co-localization analysis. Plots showing representative individual signal intensity and distribution of CDH2, PAPC and F-Actin (Phalloidin) on confocal sections along an individual cell-cell junction in the rostral half of Somite 0 (B) or Somite I (C). Note the absence of PAPC in the SI domain. These intensity profiles where used to cross-correlate the signal using IgorPro software as quantified in Fig. 3G (see Material and Methods for details).

**(D-E)** Cell-cell junction co-localization analysis. Plots showing representative individual cross-correlation signals obtained from the Somite 0 rostral half and Somite I (in D and E) respectively . The pearson coefficient is represented over the junction length indicating how much and where the signals are the most similar. A maximum at 0μm means that most of the signal in both channels co-localize, at a 200nm resolution.

**Figure S3**

**(A)** Expression of *PAPC* in pRFP control (upper panel) and PAPC-RNAi (lower panel) electroporated two-day-old chicken embryos. *PAPC* mRNA is detected by *in situ* hybridization. Arrowheads mark the last formed boundary. Dorsal view, anterior to the top. Scale bar, 200μm.

**(B)** Expression of PAPC protein (green) in the anterior PSM electroporated with the control vector pRFP or PAPC-RNAi constructs (red). Nuclei are stained with Dapi (blue). Scale bar, 10μm.

**Figure S4**

**(A)** Representative confocal images of the cellular localization of GFP (green), PAPC (magenta), and Dextran (red) in embryos electroporated with pCImG control vector or PAPC-S construct and subsequently treated with either DMSO (left) or the clathrin-mediated endocytosis inhibitor Pitstop2 (right). Regions of interest with electroporated GFP+ cells are delimited by a solid white line. Scale bar, 15μm.

(B) Quantification of Dextran uptake as a measure of endocytosis level. Signal intensity of fluorescent dextran was measured in cells treated as described in (A). Dextran signal intensity in GFP+ and GFP-cells in pCImG or PAPC-S electroporated regions and subsequently treated with DMSO (control) or Pitstop2 (red). Mean +/− s.d. t-test p values are indicated.

**Figure S5**

**(A-D)** CDH2 and ZO-1 distribution by immunofluorescence in chicken PSM explants cultured 3 hours in the presence of DMSO (0.2%) (A,C) and Chlorpromazine (50μM) (B,D). Dorsal view, anterior to the top. t0: forming somite at the time of treatment start. Somite limits are highlighted by dashed white lines. Yellow arrowheads show posterior epithelial wall assembly. Dorsal views, anterior to the top. Scale bar, 100μm.

